# Lateral entorhinal cortex afferents reconfigure the activity in piriform cortex circuits

**DOI:** 10.1101/2024.06.16.599205

**Authors:** Olivia Pedroncini, Noel Federman, Antonia Marin-Burgin

## Abstract

Odours are key signals for guiding spatial behaviours such as foraging and navigation in rodents. It has recently been found that odour representations in the piriform cortex (PCx) can also contain information about their spatial context. However, the precise origins of this information within the brain and its subsequent integration into the microcircuitry of the PCx remains unknown. In this study, we focus on the lateral entorhinal cortex (LEC) as a candidate for carrying spatial contextual information to the PCx, to investigate how it affects the PCx microcircuit and its response to olfactory inputs. Utilising mice brain slices, we performed patch clamp recordings targeting both superficial (SP) and deep (DP) pyramidal neurons, as well as parvalbumin (PV) and somatostatin (SOM) inhibitory interneurons. Concurrently, we optogenetically stimulated excitatory LEC projections to study their impact on PCx activity. We found that LEC inputs are heterogeneously distributed in the PCx microcircuit, evoking larger excitatory currents in SP and PV neurons compared to DP and SOM neurons, respectively, due to their higher monosynaptic connectivity. Moreover, LEC inputs exert a differential effect on the inhibitory circuits, activating PV while suppressing SOM interneurons. We further studied the interaction among LEC inputs and the sensory afferent signals originating from the lateral olfactory tract (LOT) onto the PCx. Our findings demonstrated that both SP and DP neurons show a general increase in their spiking response when LEC and LOT are simultaneously activated. Notably, DP neurons exhibit a sharpening of their response attributable to LEC-induced inhibition that effectively suppresses the delayed spikes evoked by LOT stimulation. These observations suggest a regulatory mechanism whereby LEC inputs inhibit recurrent activity by activating PV interneurons. Our results show that LEC afferents reconfigure PCx activity and contribute to the understanding of how odour objects are formed within the PCx integrating both olfactory and contextual information.

**Significant statement:** Primary sensory cortices are more complex than initially thought, encoding movement-related information, as well as spatial maps of animal location. The primary olfactory cortex, the piriform cortex (PCx), not only responds to the presence of odours, but to other non-olfactory signals like the spatial context in which odours are presented. The source of the contextual modulation is not known. In this work, we studied the modulation of PCx neurons by afferents arriving from the lateral entorhinal cortex (LEC), an area related to spatial representation. By the use of optogenetic to activate LEC inputs we found that its activation recruits excitatory and inhibitory circuits in PCx. The interaction of LEC inputs with afferents carrying odour information arriving from the lateral olfactory tract (LOT) resulted in a sharpening PCx principal neurons response to LOT, by a reduction in recurrent activity through PV interneurons. This circuital reorganisation of activity in PCx by LEC afferents could be an important mechanism to switch responses to be favouring dominance of the LOT pathway over the recurrent pathway. These results shed light on the understanding of how contextual aspects of animal experience, in which LEC is involved, can influence odour processing potentially leading to a richer representation of odour objects.

## Introduction

The complex and fluctuating nature of the environment in which we live requires the nervous system to be capable of generating flexible and adaptable internal representations of sensory stimuli. Previous works have demonstrated that this flexibility occurs early at the level of primary cortices, modulating the representation of a given stimulus depending on the task requirements (Poort et al., 2015). Recent studies on the olfactory processing of rodents have revealed the presence of spatial representations within the olfactory cortex when odours are associated with locations (Poo et al., 2022). Furthermore, it has been demonstrated that spatial and other non-olfactory information emerge in the olfactory cortex after learning (Federman et al., 2023).

The piriform cortex (PCx) is the largest region of the olfactory cortex and plays a crucial role in encoding odour objects (Gottfried, 2010; D. A. Wilson & Sullivan, 2011). It is a trilaminar cortex and comprises two distinct populations of pyramidal neurons: the superficial pyramidal (SP) neurons located in layer 2 and deep pyramidal (DP) neurons, in layer 3. Both populations receive direct sensory afferents from the olfactory bulb through the lateral olfactory tract (LOT). These dynamically interact with extensive recurrent or associational (ASSN) connections made by the PCx principal neurons (Franks et al., 2011; Poo & Isaacson, 2009, 2011). The proportion of inputs differs such that SP neurons are dominated by sensory afferents while DP neurons are mostly driven by recurrent circuits (Wiegand et al., 2011).

Moreover, afferent inputs not only excite PCx neurons but also recruit inhibitory circuits in a feedforward and feedback manner (Franks et al., 2011; Stokes & Isaacson, 2010). The feedback inhibition is primarily mediated by Parvalbumin (PV) and Somatostatin (SOM) interneurons (Suzuki & Bekkers, 2010). PV interneurons make perisomatic synapses with the principal neurons (Stokes & Isaacson, 2010; Suzuki & Bekkers, 2010) and are thought to regulate their spike activity (Canto-Bustos et al., 2022). Conversely, SOM interneurons establish synapses on the proximal apical dendrites of the principal neurons influencing dendritic integration of afferent and recurrent inputs (Large, Kunz, et al., 2016). Therefore, it’s the interplay between excitatory and inhibitory circuits in the PCx that determines the population of neurons activated in response to a stimulus.

In addition to the LOT and ASSN inputs, the PCx is reciprocally and extensively interconnected with higher-order cortical areas associated with emotional learning and mnemonic processes, such as the orbitofrontal cortex (Illig, 2005), the amygdala (Majak et al., 2004), and the perirhinal and entorhinal cortices (Chapuis et al., 2013; Johnson et al., 2000). All these regions send excitatory feedback projections to the PCx that may contribute to the formation of a broader and experience-dependent sensory representation in the PCx. One of these regions, the lateral entorhinal cortex (LEC), has been largely studied due to its strong connection with the hippocampus and has been associated with the processing of the spatial location of objects. The interaction of LEC afferents with the PCx circuit could confer spatial modulation to the olfactory representation observed in the activity of PCx neurons when animals learn to associate odours with spatial contexts and rewards (Federman et al., 2023; Poo et al., 2022). However, the underlying mechanism remains unknown.

In this study we aimed to understand how the projections of excitatory LEC neurons are integrated into the PCx microcircuit exploring the response of different subpopulations of PCx neurons to the activation of this input. We found that LEC afferents establish monosynaptic connections with SP neurons, and its activation leads to an enhancement of SP response to LOT stimulation. Notably, DP neurons exhibited a marked suppression of their late response to LOT stimulation which could be a consequence of the ASSN pathway silencing. Our findings suggest that the reduction in recurrence may be attributed to the activation of inhibitory circuits, with PV interneurons playing a central role. This could be a crucial mechanism for switching responses to favour the dominance of the LOT pathway over the recurrent pathway. Overall, these results highlight how contextual aspects of animal experience, involving the LEC, can influence odour processing, potentially leading to a richer representation of odour objects.

## Methods

### Subject details

To generate *Pvalb^Cre^*;CAG^FloxStopTom^ (PV-Cre-floxTom) mice, *B6;129P2-Pvalbtm1(cre)Arbr/J (PV^Cre^)* mice (Hippenmeyer et al., 2005) were crossed with *B6.Cg-Gt(ROSA)26Sortm14(CAG-tdTomato)Hze/J (Ai14)* conditional reporter mice. Cre heterozygous animals were used.

To generate *SOM^Cre^*;CAG^FloxStopTom^ (SOM-Cre-floxTom) mice, *Ssttm2.1(cre)Zjh/J (SOM^Cre^)* were crossed with *B6.Cg-Gt(ROSA)26Sortm14(CAG-tdTomato)Hze/J (Ai14)* conditional reporter mice. Cre heterozygous animals were used.

Mice used in this study were both male and female, and aged from 4 to 11 weeks (see below for specific experiments). Mice were housed under controlled environment in a 12h light/dark cycle, with food and water *ad libitum*.

Experimental protocol (2020-03-NE) was evaluated by the Institutional Animal Care and Use Committee of the IBioBA-CONICET according to the Principles for Biomedical Research involving animals of the Council for International Organizations for Medical Sciences and provisions stated in the Guide for the Care and Use of Laboratory Animals.

### Animals and surgery for virus delivery

For SP and DP recordings, we employed male C57BL/6J mice. For interneurons recordings, we use either sex PV-Cre-floxTom transgenic mice, and either sex SOM-Cre-floxTom transgenic mice. All experimental mice were aged between 4 to 6 weeks and weighed over 15 grams at the time of surgical procedures.

For surgery, mice were anaesthetised (150 μg ketamine/15 μg xylazine in 10 μl saline/g), and 400-450nl of adenoviral vector pAAV-CaMKIIa-hChR2(H134R)-mCherry (*AddGene)* was infused at 125 nl/min into the left LEC using sterile microcapillary calibrated pipettes (Sigma) and stereotaxic references (coordinates from bregma: −3.6 mm anteroposterior,+3.9 mm mediolateral, −3 mm dorsoventral).

### Slice preparation

After a 4-5 weeks post-viral injection period to allow Chr2 expression along infected neuron projections, mice were anaesthetised and euthanized by decapitation. Brains were removed into a chilled solution containing (mM) 110 choline-Cl^−^, 2.5 KCl, 2.0 NaH_2_PO_4_, 25 NaHCO_3_, 0.5 CaCl_2_, 7 MgCl_2_, 20 dextrose, 1.3 Na^+^-ascorbate, 3.1 Na^+^-pyruvate. The left hemisphere was removed and slices (350 μm thick) were cut coronally in a vibratome, and transferred to a chamber containing artificial cerebrospinal fluid (ACSF; mM): 125 NaCl, 2.5 KCl, 2.3 NaH_2_PO_4_, 25 NaHCO_3_, 2 CaCl_2_, 1.3 MgCl_2_, 1.3 Na^+^-ascorbate, 3.1 Na^+^-pyruvate, and 10 dextrose (315 mOsm). Slices were bubbled with 95% O_2_/5% CO_2_ and maintained at 30°C for approximately 1 hr before experiments started. Salts were acquired from Sigma-Aldrich (St. Louis, MO). When tetrodotoxin (TTX, 1 μM, Tocris) and 4-aminopyridine (4-AP, 100 μM, Sigma Aldrich) were used, these were perfused into the bath solution.

### Electrophysiological recordings

SP and DP recorded neurons were identified by their layer location (SP located in packed layer 2, DP in more sparsed layer 3) using infrared DIC video microscopy. PV and SOM interneurons expressing fluorescent protein dTomato were identified using a green LED source delivered through the epifluorescence pathway of the upright microscope.

Whole-cell recordings were performed using microelectrodes (4–10 MΩ) filled with (in mM) 130 CsOH, 130 D-gluconic acid, 2 MgCl2, 0.2 EGTA, 5 NaCl, 10 HEPES, 4 ATP-tris, 0.3 GTP-tris, 10 phosphocreatine. In experiments where spiking responses were evaluated, a potassium gluconate internal solution was used (in mM): 120 potassium gluconate, 4 MgCl2, 10 HEPES buffer, 0.1 EGTA,, 5 NaCl, 20 KCl, 4 ATP-tris, 0.3 GTP-tris, and 10 phosphocreatine (pH = 7.3; 290 mOsm).

Recordings were obtained using Multiclamp 700B amplifiers, (Molecular Devices), digitised, and acquired at 20 KHz onto a personal computer using the pClamp10 software. Membrane capacitance and input resistance were obtained from current traces evoked by a hyperpolarizing step of 10 mV. Series resistance was typically 10– 20 MΩ, and experiments were discarded if higher than 40 MΩ.

### Evoked postsynaptic currents

Evoked monosynaptic excitatory postsynaptic currents (EPSC) and inhibitory postsynaptic currents (IPSC) were recorded after optogenetic stimulation executed by a 470 nm LED source delivered through the epifluorescence pathway of the upright microscope and commanded by the acquisition software. The stimulation protocol consisted of trains of 5 blue light pulses (10Hz, pulse width=0.5 ms, maximum intensity) delivered every 30 seconds.

EPSCs were isolated by voltage clamping SP, DP, PV or SOM at the reversal potential of the IPSC measured for each individual neuron (∼-70 mV). In turn, IPSCs were recorded at the reversal potential of the EPSC (∼0 mV). When possible, both EPSC and IPSC were recorded from the same cell.

Charge was calculated as the integral of the current over a 90 ms window following the pulse. Excitatory current with peaks below −800pA were considered spikes and were excluded from EPSC analysis. For latency analysis, recordings with average current peaks below 20pA were excluded to ensure accurate onset determination.

The excitatory to inhibitory balance was calculated as the proportion of excitatory charge over the total charge (sum of excitatory and inhibitory charge). This way of expressing the balance is more convenient than a simple ratio when there are values close to zero in the sample, as in our case.

### Monosynaptic responses

In order to dissect LEC monosynaptic inputs a combination of Tetrodotoxin (1 µM) and 4-aminopyridine (100µM) was added to the perfusion bath. TTX blocks voltage-gated sodium channels, preventing the generation of action potentials (APs), while 4-AP blocks potassium channels, depolarizing cell membranes and facilitating the release of neurotransmitter vesicles from stimulated terminals.

At least 5 minutes were allowed to pass between the addition of the drugs and the assessment of their effect. A threshold of −15pA was established to determine whether a neuron responded or not under the influence of the TTX and 4-AP drugs.

### Spiking Response

Single APs were identified when the membrane voltage exceeded a threshold set at V=50mV. To study LOT responses, a stimulation protocol comprising 5-pulse trains (10Hz, pulse width==0.2ms, threshold intensity) delivered by an electrode in the LOT was applied every 30 seconds. To study responses to both LOT and LEC pathways, this protocol was combined with optogenetic stimulation (5-pulse trains at 10hz, pulse width=0.5ms, maximum intensity) aligning the onset of electrical and light pulses. A search window of 70ms was established following each stimulation pulse for AP detection. Latency was calculated as the time between the peak of the pulse and the peak of the action potential. APs with latencies smaller than 2ms were excluded from the analysis, as they were considered to result from direct rather than synaptic stimulation.

### Quantification and statistical analysis

Unless otherwise specified, data is presented as mean ± SEM. Normality was assessed using Shapiro-Wilk’s test, D’Agostino & Pearson omnibus test, or Kolmogórov-Smirnov’s test, with a *p* value of 0.05. Two-tailed ordinary, one sample or paired t test was used for single comparisons, and ordinary or repeated-measures ANOVA for multiple comparisons, with post hoc Tukey’s test. For non-normal distributions, comparisons were made using non-parametric tests such as Mann-Whitney test and Wilcoxon signed rank test. Statistical details of experiments can be found in the figure legends and results section.

## Results

### LEC afferents are heterogeneously distributed in the PCx microcircuit

In order to explore the effect of LEC activation in the piriform microcircuit, we injected an adeno-associated virus carrying the photoactivatable protein channelrhodopsin-2 (ChR2) under a promoter of excitatory neurons (Camk2a) in LEC. We then obtained ipsilateral PCx brain slices and stimulated them with blue light (Fig. 1A), while performing electrophysiological whole-cell recordings from different groups of excitatory and inhibitory neurons in PCx (Fig. 1B). We first recorded, in voltage-clamp mode, excitatory postsynaptic currents (EPSC) from the two populations of pyramidal cells, SP and DP neurons, in response to LEC stimulation with a train of 5 pulses at a frequency of 10Hz (Fig. 1C). We observed LEC-evoked EPSCs in 62.8% of the recorded SP neurons (n=44 cells) and in 74.2% of the DP neurons (n=31 cells) (Fig. 1 D). Comparing those responsive neurons, we found that SP neurons receive significantly larger currents than DP neurons along the stimulation train (two-way ANOVA, SP versus DP: F = 4.591, p=0.0376; variation in pulse number: F= 8.002, p = 0.0008, interaction: F= 0.0998, p=0.9824; n_SP_=26 cells, n_DP_=21 cells) (Fig. 1E). However, when we compared the total charge evoked by the sum of the five pulses of the train, we observed a non-significant trend (Mann Whitney test, p=0.0975, n_SP_=26 cells, n_DP_=21 cells) that, we believe, is due to a big variability within each group (Fig. 1F).

**Figure 1:**
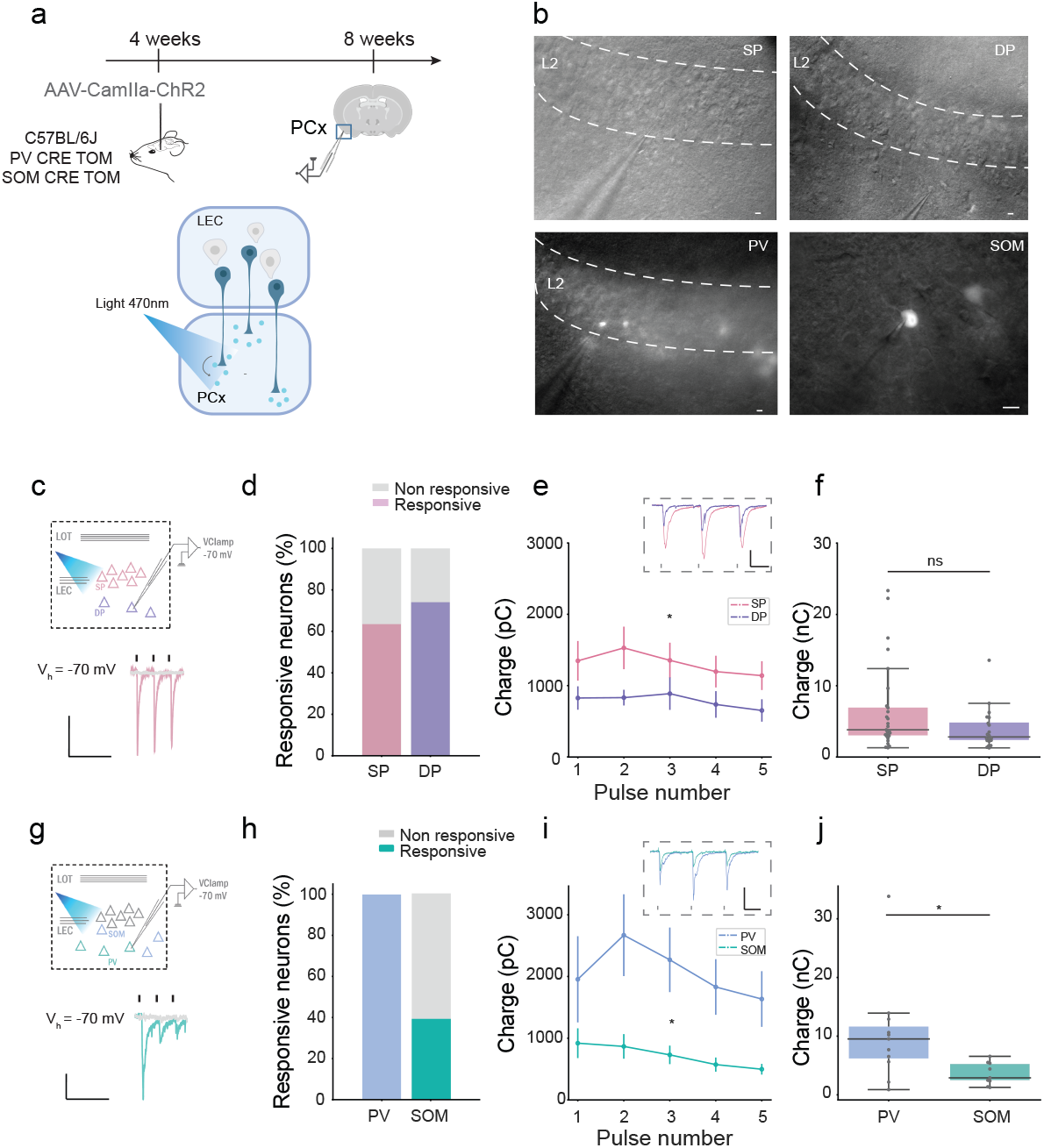
Distinct effect of LEC input among PCx neuronal populations. **A,** Experimental design. TOP: Experimental timeline. 4-week animals are injected with the adenovirus expressing ChR2; after 4 weeks of viral expression, animals are sacrificed, and acute brain slices of piriform cortex are preserved for electrophysiological recordings. BOTTOM: Optogenetic strategy. Blue light over piriform cortex slice promotes neurotransmitter release from terminals of neurons expressing ChR2. **B,** Representative images of electrophysiological recordings from Superficial Pyramidal neurons (SP), Deep Pyramidal neurons (DP), Parvalbumin Interneurons (PV) and Somatostatin interneurons (SOM). Scale bar: 20µm. **C,D,E,F,** Whole-cell recordings of EPSCs evoked by LEC stimulation in pyramidal neurons. **C,** TOP: Scheme illustrating experimental procedure. Individual SP and DP neurons are recorded in voltage-clamp holding the membrane at Vh = −70mV while stimulating with blue light delivered at 10Hz in a 5-pulses train. BOTTOM: Representative traces of a responsive neuron (pink) and a non-responsive neuron (grey). Black rectangles represent each stimulation pulse (only the first 3 pulses are shown). Scale bar: x=250ms, y=100pA. **D,** Percentage of SP and DP responsive neurons. **E,** Charge of EPSC evoked by each pulse of LEC stimulation train in SP and DP responsive neurons (two-way ANOVA, SP versus DP: F = 4.591, p=0.0376; variation in pulse number: F= 8.002, p = 0.0008, interaction: F= 0.0998, p=0.9824; n_SP_=26 cells, n_DP_=21 cells). INSET: Representative traces of a SP and a DP EPSC. Scale bar: x=50ms, y=100pA. **F,** Total charge of EPSCs evoked by the stimulation train (Mann Whitney test, p=0.0975, n_SP_=26 cells, n_DP_=21 cells). **G,H,I,J,** Whole-cell recordings of EPSCs evoked by LEC stimulation in inhibitory interneurons. **G,** TOP: Scheme illustrating experimental procedure. Individual PV and SOM interneurons are recorded in voltage-clamp holding the membrane at Vh = −70mV and stimulated with blue light with a 10Hz 5-pulses train. BOTTOM: Representative traces of a responsive neuron (green) and a non-responsive neuron (grey). Black rectangles represent each stimulation pulse (only the first 3 pulses are shown). Scale bar: x=250ms, y=20pA. **H,** Percentage of PV and SOM responsive neurons. **I,** Charge of EPSC evoked by each pulse of LEC stimulation train in PV and SOM responsive neurons (two-way ANOVA, PV versus SOM: F=5.8, p= 0.0367; variation in pulse number: F= 5.095, p= 0.0084; interaction: F=1.918, p=0.1166;, nPV=11 cells, nSOM=9 cells). INSET: Representative traces of a PV and a SOM EPSC. Scale bar: x=50ms, y=100pA. **J,** Total charge of EPSCs evoked by the stimulation train (Mann Whitney test: p=0.0159, nPV=11 cells, nSOM=9 cells).

Next, we studied the interaction of LEC inputs with inhibitory circuits of PCx. We recorded LEC-evoked EPSCs from two of the main populations of GABAergic neurons present in PCx, PV and SOM interneurons. For these experiments, we used transgenic mouse lines PV-Cre-floxTom and SOM-Cre-floxTom that allowed us to identify PV-IN and SOM-IN respectively (Fig. 1.B). We detected excitatory responses in 100% of PV and only 39.1% of SOM interneurons (n_PV_=11 cells, n_SOM_=23 cells) (Fig. 1.G,H). Taking the subset of responsive neurons, we analysed EPSCs evoked by a 5 pulses train stimulation of blue light, and observed that the currents in PV interneurons were larger than those in SOM (two-way ANOVA, PV versus SOM: F=5.8, p= 0.0367; variation in pulse number: F= 5.095, p= 0.0084; interaction: F=1.918, p=0.1166;, nPV=11 cells, nSOM=9 cells) (Fig. 1I). We obtained the same result comparing the total charge of the stimulation train (Mann Whitney test: p=0.0159, nPV=11 cells, nSOM=9 cells) (Fig. 1J).

### LEC projections preferentially contact SP neurons and PV interneurons

As PCx neurons are extensively interconnected forming a recurrent network, we were uncertain whether the EPSCs that we recorded were due to direct monosynaptic input from LEC or, alternatively, recurrent excitation. To address this, we quantified the latency of EPSCs for each neuronal population (Fig. 2.A). Both SP and PV neurons showed similar onset latencies (median_SP_ = 4.983 ± 1.51 ms, n = 119 events; median_PV_=5.16 ± 3.339 ms, n=60 events). SP latencies were significantly shorter than the ones of DP and SOM neurons (median_DP_ =6.873 ± 3.234, median_SOM_ =5.418 ± 4.324; Kruskal-Wallis test, statistic=28.82, p<0.0001; Dunn’s multiple comparison, SP versus DP: mean rank diff=-62.55, p<0.0001; SP versus PV: mean rank diff=-35.96, p= 0.075; SP versus SOM: mean rank diff=-57.77, p =0.006; n_SP_=119 events, n_DP_=102 events, n_PV_=60 events, n_SOM_=35 events) (Fig. 2B). The difference in the latency to evoke a postsynaptic current could reflect variations in the connectivity of LEC inputs among each neuronal population. These results suggest that the recorded response in DP and SOM neurons should be largely attributed to the recurrent activity of the circuit rather than to direct stimulation from LEC.

**Figure 2:**
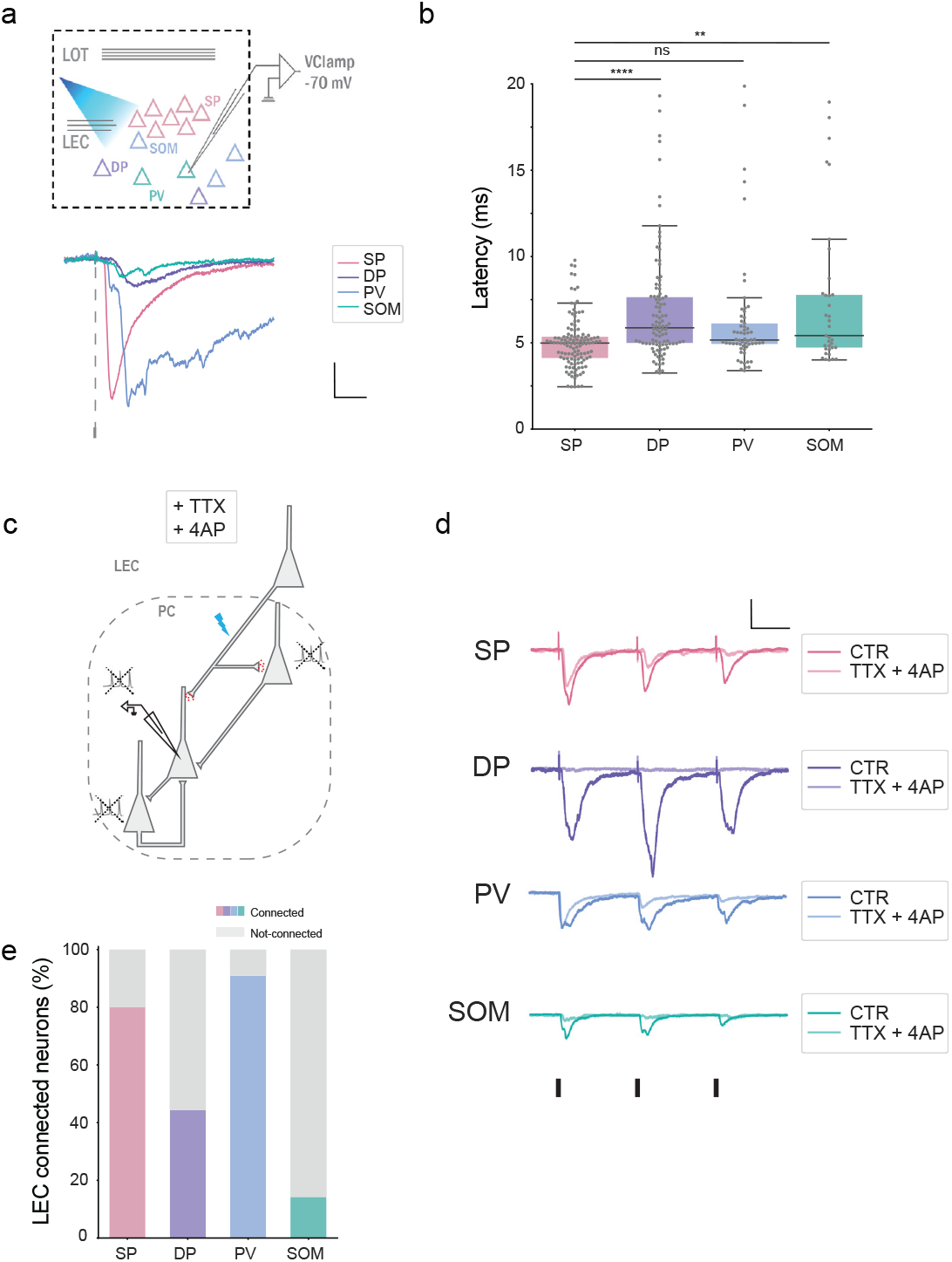
Excitatory LEC projection contacts preferentially SP and PV neurons in the PCx. **A,** TOP: Scheme illustrating experimental procedure. Individual SP, DP, PV and SOM neurons are recorded in voltage-clamp holding the membrane at Vh=-70mV while stimulating with blue light delivered at 10Hz in a 5-pulses train. BOTTOM: Representative traces showing a difference in latencies of the EPSC (only first pulse is shown). Scale bar: x=10ms, y=50pA. **B**, Latencies to the onset of EPSC for different neuronal populations. Each point corresponds to an EPSC evoked by a single pulse of the train (all pulses are pooled). Boxes indicate median and quartiles (Kruskal-Wallis test, statistic=28.82, p<0.0001; Dunn’s multiple comparison, SP versus DP: mean rank diff=-62.55, p<0.0001; SP versus PV: mean rank diff=-35.96, p=0.075; SP versus SOM: mean rank diff=-57.77, p=0.006; n_SP_=119 events, n_DP_=102 events, n_PV_=60 events, n_SOM_=35 events). **C,** Pharmacological strategy to dissect monosynaptic connections. The combination of Tetrodotoxin (TTX) and 4-aminopyridine (4-AP) in the bath prevents neurons from firing action potentials, effectively maintaining membrane depolarization and thereby facilitating neurotransmitter release from stimulated terminals. **D,** Representative traces of EPSCs from SP, DP, PV and SOM neurons before (CTR) and after (TTX+4AP) the infusion of the drugs into the bath. Scale bar: x=50ms, y=50pA. **E,** Percentage of connected neurons (SP=80%, DP=44.44%, PV=90.9%, SOM=14.3%; n_SP_=15 cells, n_DP_=9 cells, n_PV_=11 cells, n_SOM_=7 cells).

In order to further clarify the connectivity of LEC projections, we recorded evoked EPSC currents in presence of Tetrodotoxin (TTX) and 4-aminopyridine (4-AP), a combination of drugs that allows dissecting monosynaptic inputs by suppressing AP firing while promoting neurotransmitter release from depolarized terminals (Fig. 2.C). We recorded LEC evoked EPSCs from SP, DP, PV and SOM neurons after the infusion of TTX and 4-AP, and classified each one as “connected” or “not-connected” by establishing a minimal response threshold (EPSCamplitude = −15pA). As expected, a high percentage of SP and PV neuronal populations were connected (80% and 90.9% respectively); whereas only a 44.4% of DP neurons and a 14.3% of SOM interneurons showed monosynaptic LEC connections (n_SP_=15 cells, n_DP_=9 cells, n_PV_=11 cells, n_SOM_=7 cells) (Fig. 2.D,E). These results demonstrate a heterogeneous connectivity of LEC afferents in PCx and suggest a modulation of activity through PV interneurons.

### Activation of LEC afferents induces a functional reorganisation of PCx inhibition

Since PV inhibitory interneurons receive direct excitatory synapses from LEC, we asked whether LEC activation induces inhibitory postsynaptic currents (IPSCs) across distinct neuronal populations within the circuit (Fig 3.A, H).

**Figure 3:**
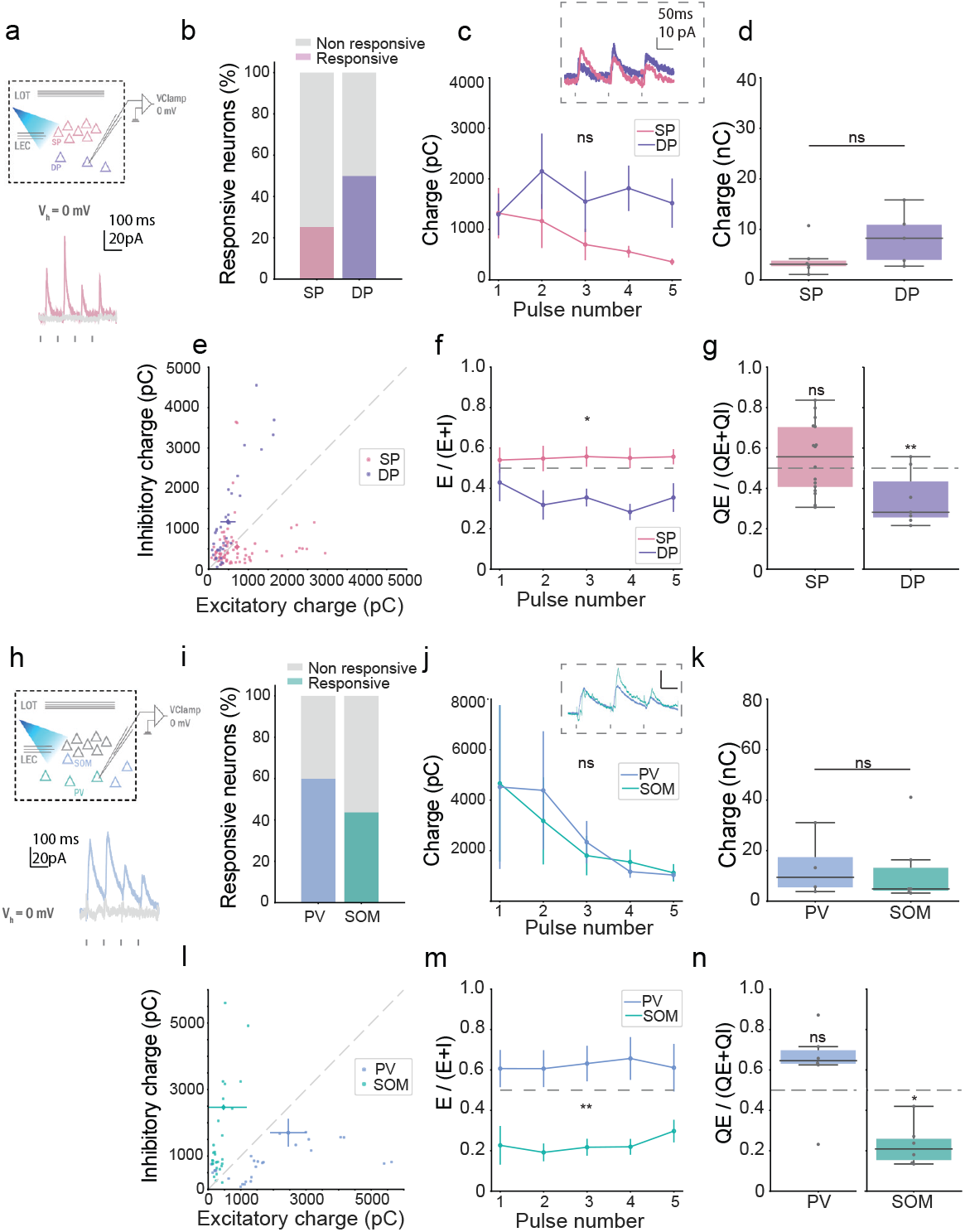
Activation of LEC input recruits inhibitory circuits in PCx. **A,B,C,** Whole-cell recordings of IPSCs evoked by LEC stimulation in pyramidal neurons. **A,** TOP: Scheme illustrating experimental procedure. Individual SP and DP neurons are recorded in voltage-clamp holding the membrane at Vh = 0mV while stimulating with blue light delivered at 10Hz in a 5-pulses train. BOTTOM: Representative traces of a responsive neuron (pink) and a non-responsive neuron (grey). Black rectangles represent each stimulation pulse (only the first 3 pulses are shown). Scale bar: x=100ms, y=20pA. **B,** Percentage of SP and DP responsive neurons. **C,** Charge of IPSC evoked by each pulse of LEC stimulation train in SP and DP responsive neurons. INSET: Representative traces of a SP and a DP IPSC. Scale bar: x=50ms, y=10pA (two-way ANOVA, SP versus DP: F=1.78, p= 0.2181; variation in pulse number: F= 1.938, p=0.1713; interaction: F=1.871, p=0.1397; n_SP_=5 cells, n_DP_=5 cells). **D,** Total charge of IPSCs evoked by the stimulation train (t-test, SP versus DP: t=1.337, p=0.2181; n_SP_=5 cells, n_DP_=5 cells). **E,F,G,** Excitation to inhibition balance for pyramidal neurons. **E,** Pairs of excitatory and inhibitory charge (all pulses are pooled). Highlighted dots illustrate mean and SEM for each neuronal population. **F,** E/I balance calculated as excitatory charge over the sum of excitatory and inhibitory charge evoked by LEC for each pulse of the train (two-way ANOVA, SP versus DP: F=7.27, p=0.0135; variation in pulse number: F= 1.053, p=0.3748; interaction: F=1.309, p=0.2735; n_SP_=16 cells, n_DP_=7 cells). **G,** Mean E/I balance for SP and DP neurons (t-test, SP versus 0.5: t=1.642, p=0.1215, n=16 cells; DP versus 0.5: t=4.956, p=0.0043, n=7 cells). **H,I,J,** Whole-cell recordings of IPSCs evoked by LEC stimulation in inhibitory interneurons. **H,** TOP: Scheme illustrating experimental procedure. Individual PV and SOM interneurons are recorded in voltage-clamp holding the membrane at Vh=0mV while stimulating with blue light delivered at 10Hz in a 5-pulses train. BOTTOM: Representative traces of a responsive neuron (blue) and a non-responsive neuron (grey). Black rectangles represent each stimulation pulse (only the first 3 pulses are shown). Scale bar: x=100ms, y=20pA. **I,** Percentage of PV and SOM responsive neurons. **J,** Charge of IPSC evoked by each pulse of LEC stimulation train in PV and SOM responsive neurons. INSET: Representative traces of a PV and a SOM IPSC. Scale bar: x=50ms, y=100pA (two-way ANOVA, PV versus SOM: F=0.0153, p=0.9044; variation in pulse number: F= 2.641, p=0.1385; interaction: F=0.1159, p=0.976; n_PV_=4 cells, n_SOM_=6 cells). **K,** Total charge of IPSCs evoked by the stimulation train (Mann-Whitney test, p=0.6095). **L,M,N** Excitation to inhibition balance for inhibitory neurons. **L,** Pairs of excitatory and inhibitory charge (all pulses are pooled). Highlighted dots illustrate mean and SEM for each neuronal population. **M,** E/I balance calculated as excitatory charge over the sum of excitatory and inhibitory charge evoked by LEC for each pulse of the train (two-way ANOVA, PV versus SOM: F=16.33, p=0.0024; variation in pulse number: F=0.458, p=0.5953; interaction: F=0.6095, p=0.6581; n_PV_=6 cells, n_SOM_=6 cells). **N,** Mean E/I balance for PV and SOM neurons (Wilcoxon signed rank test, PV vs. 0.5: W=11, p=0.3125, n=6 cells; SOM versus 0.5: W=-21, p= 0.0313, n=6 cells).

We observed LEC-evoked IPSCs in 25% of the SP neurons we recorded (n_SP_=20 cells) and in 50% of the DP neurons (n_DP_=10 cells) (Fig 3.B).We did not find any significant difference between them when we compared the inhibitory charge per pulse (two-way ANOVA, SP versus DP: F=1.78, p= 0.2181; variation in pulse number: F= 1.938, p=0.1713; interaction: F=1.871, p=0.1397; n_SP_=5 cells, n_DP_=5 cells) or the total charge (t-test, SP versus DP: t=1.337, p=0.2181; n_SP_=5 cells, n_DP_=5 cells) (Fig. 3.C, D).

To understand how individual neurons integrate the inputs resulting from LEC activation, we evaluated the excitation to inhibition balance (E/I balance) calculated as the amount of excitation over the sum of excitation and inhibition (E/E+I). We considered all the neurons that we recorded at both V_h_=-70mV and V_h_=0mV (n_SP_=16 cells, n_DP_=7 cells) (Fig.3.E). Neurons that exhibited neither EPSCs nor IPSCs in response to LEC activation were excluded from the analysis. The two groups of pyramidal neurons showed a difference in the E/I balance along the train (two-way ANOVA, SP versus DP: F=7.27, p=0.0135; variation in pulse number: F= 1.053, p=0.3748; interaction: F=1.309, p=0.2735; n_SP_=16 cells, n_DP_=7 cells) (Fig. 3.F). For SP neurons, the E/I balance was close to 0.5 indicating that a given magnitude of excitatory current elicits a correspondingly similar inhibitory current (t-test, SP versus 0.5: t=1.642, p=0.1215, n=16 cells). In contrast, for DP neurons, inhibitory charge overcame excitatory charge leading to a “negative” E/I balance (t-test, DP versus 0.5: t=4.956, p=0.0043, n=7 cells) (Fig. 3.G). This suggests that activation of LEC inputs would result in an overall activation of SP and an inhibition of DP neurons.

In addition, we recorded IPSCs and calculated E/I balance for PV and SOM interneurons (Fig 3.H). We did not find any difference in the inhibitory charge per pulse (two-way ANOVA, PV versus SOM: F=0.0153, p=0.9044; variation in pulse number: F= 2.641, p=0.1385; interaction: F=0.1159, p=0.976; n_PV_=4 cells, n_SOM_=6 cells) or the total charge (Mann Whitney test, PV versus SOM: p=0.6095) (Fig. 3.L,M,N). Consistent with the difference in EPSCs previously reported, the resulting E/I balance per pulse differed between PV and SOM interneurons (two-way ANOVA, PV versus SOM: F=16.33, p=0.0024; variation in pulse number: F= 0.458, p=0.5953; interaction: F=0.6095, p=0.6581; n_PV_=6 cells, n_SOM_=6 cells) (Fig 3.M). SOM interneurons showed a striking “negative” (under 0.5) mean E/I balance (t-test, SOM versus 0.5: t=6.157, p= 0.0016, n=6 cells) meaning that inhibitory inputs predominate over excitatory ones. In the case of PV interneurons, even though we did not observe a significant E/I balance above 0.5, we detected spiking activity in response to light (unlike the rest of the neuronal populations recorded) (Fig S1). These results demonstrate that inputs arriving from LEC have a different impact in both populations of interneurons, which would lead to a reorganisation of the PCx output.

### LEC afferent activation modulates pyramidal spiking responses to LOT inputs

The effect of stimulating the isolated LEC pathway resulted in a differential activation of different populations of neurons in PCx. We then wanted to evaluate whether this input could modulate how pyramidal neurons respond to afferent input arriving from the olfactory bulb. To study this, we recorded the spiking activity of SP and DP neurons in response to either LOT electrical stimulation, or combined LOT and LEC simultaneous stimulation. We set the intensity of the stimulation electrode placed in the LOT at threshold level, and stimulated with a train of 5 pulses delivered at 10 Hz. For combined stimulation of LOT and LEC, we superimposed a 5 pulses train at 10Hz of blue light at a fixed intensity, to the LOT stimulation protocol (Fig 4.A).

**Figure 4:**
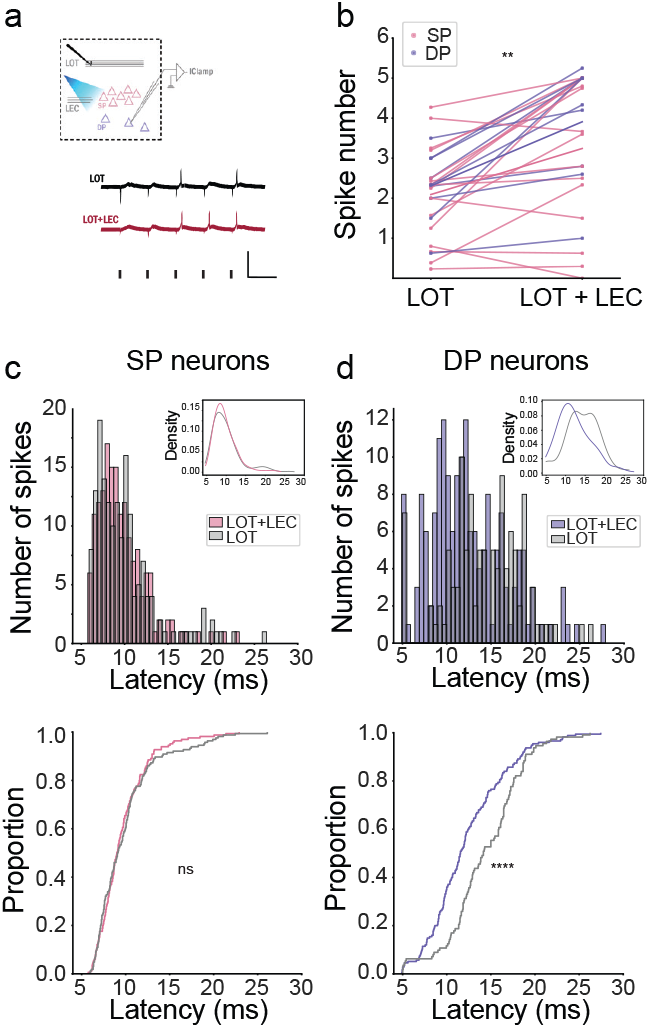
LEC activation modulates pyramidal spiking responses to OB afferent input. **A,** TOP: Scheme illustrating experimental procedure. Individual SP and DP neurons are recorded in current-clamp mode. LOT is stimulated electrically with a 5-pulses train delivered at 10Hz frequency and threshold intensity. LEC is stimulated with blue light in a 5-pulses train at 10Hz. BOTTOM: Representative traces showing spiking in response to either LOT stimulation or simultaneous LOT and LEC stimulation. Scale bar: x=100ms, y=1V. **B,** Total number of action potentials in response to LOT train stimulation or combined LOT and LEC simultaneous stimulation (paired t-test, SP_LOT_ versus SP_LOT+LEC_: t=3.374, p=0.0042, n=16 pairs; DP_LOT_ versus DP_LOT+LEC_: t=4.371, p=0.0024, n=9 pairs). **C,** TOP: Histogram showing the distribution of latencies to spike occurrence for SP neurons (all pulses are pooled). INSET: kernel density estimation for LOT (grey) and LOT+LEC (pink) distributions. BOTTOM: Cumulative distribution of latencies in response to LOT stimulation (grey) or combined LOT and LEC stimulation (pink) (Kolmogorov-Smirnov test, statistic=0.0667, p=0.8584, n_LOT_=165 spikes, n_LOT+LEC_=165 spikes). **D,** TOP: Histogram showing the distribution of latencies to spike occurrence for DP neurons (all pulses are pooled). INSET: kernel density estimation for LOT (grey) and LOT+LEC (blue) distributions. BOTTOM: Cumulative distribution of latencies in response to LOT stimulation (grey) or combined LOT and LEC stimulation (blue) (Kolmogorov-Smirnov test, statistic= 0.2803 p<0.0001, n_LOT_=112 spikes, n_LOT+LEC_=175 spikes).

We observed an increase in the number of evoked spikes when stimulating the combined pathways for both SP and DP neurons (paired t-test, SP_LOT_ versus SP_LOT+LEC_: t=3.374, p=0.0042, n=16 pairs; DP_LOT_ versus DP_LOT+LEC_: t=4.371, p=0.0024, n=9 pairs) (Fig. 4.B). However, when we analysed the temporal profile of these responses (i.e., the latency to the peak of each action potential) we noticed, only for DP neurons, a temporal shift towards earlier responses when both pathways were stimulated (Kolmogorov-Smirnov test, SP_LOT_ versus SP_LOT+LEC_: S=0.067, p=0.8584, n_LOT_=165 APs, n_LOT+LEC_=165 APs; DP_LOT_ versus DP_LOT+LEC_: S= 0.2804, p <0.0001, n_LOT_=112 APs, n_LOT+LEC_=175 APs) (Fig 4.C,D). It’s worth noting that the basal response of SP neurons to LOT stimulation exhibits an earlier and sharper distribution compared to the spiking response of DP neurons. This can be explained due to the prevailing influence of LOT input over associational one on SP neurons. The broad basal distribution of DP neurons is consistent with a larger influence of the ASSN pathway. From this result, we propose that LEC predominantly affects the recurrence of the circuit, thereby producing a more pronounced effect on DP neurons.

An increase in the total number of APs elicited and a temporal lock to the activated pathways, made us think that DP neurons could be acting as coincidence detectors of LOT and LEC inputs. To further explore this hypothesis, we designed an experiment that could reveal two key aspects: the silencing of late responses, and the sharpening of early ones. We stimulated LOT with a sequence of 2 pulses varying the intertrial interval (ITI) from 15 to 30 ms (Fig. 5.A). The different ITIs were selected based on the spiking latencies previously shown (Fig. 4.D) expecting to find a time window in which the recurrence exerted more influence in the spiking response after the second pulse. The stimulation intensity was maintained such that the second pulse reached the threshold level. We quantified the percentage of spike occurrences following the second pulse and compared this with a condition in which we superimposed a single light pulse to the first LOT pulse. We found that, when adding LEC stimulation, the percentage of spike occurrence elicited by the second LOT pulse decreased for ITI=25 ms (t-test, %Change_P2_ versus 0, t=3.419, p=0.0419, n=4 cells) (Fig. 5.B,C) implying a contraction in the summation window. Additionally, we noticed that the distribution of spikes elicited by the first pulse became sharper (SD_LOT_=3.6ms, SD_LOT+LEC_=2.17ms) (Fig.5.D). Hence, these findings support the hypothesis that DP neurons function as precise coincidence detectors locking the spiking response to the timing of afferent inputs, and further demonstrate the suppression of late spikes.

**Figure 5:**
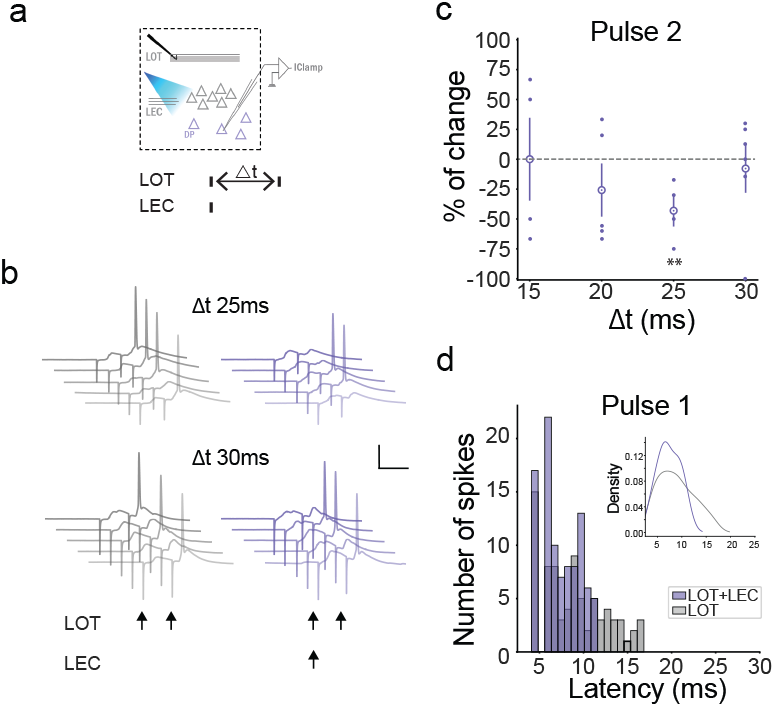
DP neurons act as coincidence detectors for LOT and LEC inputs. **A,** Scheme illustrating experimental procedure. Individual DP neurons are recorded in current-clamp mode. LOT is stimulated electrically with 2 pulses separated by a moving interval ranging from 15ms to 30ms, at threshold intensity. LEC is stimulated with a single pulse of blue light coincident with LOT first pulse. **B,** Example traces of one DP neuron spiking for two different ITIs (25ms and 30ms). Scale bar: x=25mV, y=25ms. **C,** Percentage of change in the second pulse evoked spiking when LEC stimulation is superimposed to the first LOT pulse for different LOT-ITIs ranging from 15ms to 30ms (t-test, %Change_15ms_ versus 0: t=0, p>0.9999, n=4 cells; %Change_20ms_ versus 0: t=1.195, p=0.2981, n=5 cells; %Change_25ms_ versus 0:t=3.419, p=0.0419, n=4 cells; %Change_30ms_ versus 0: t=0.3971, p=0.7077, n=6 cells). **D,** Histogram showing the distribution of latencies to first spike occurrence in response to the first pulse of stimulation. INSET: kernel density estimation for LOT (grey) and LOT+LEC (blue) distributions.

## Discussion

In this work we found that LEC inputs are heterogeneously distributed in the PCx microcircuit. Specifically, we observed larger responses in SP neurons and PV interneurons compared to DP neurons and SOM interneurons (Fig. 1.E,F). Consistent with this result, our analysis revealed a preferential connection of LEC terminals with SP and PV neurons (Fig. 2.E). Interestingly, we found that LEC inputs differentially affect the inhibitory circuits of the PCx, activating PV while suppressing SOM interneurons (Fig S1, Fig. 3.L-N). Furthermore, our experiments demonstrated that LEC activation modulates the responses of pyramidal neurons to inputs from the LOT. While both SP and DP neurons exhibited an overall increase in spiking response (Fig. 4.B), LEC activation produced a sharpening of DP neuronal activation by abolishing late components of the response to LOT (Fig. 4.D, Fig. 5.C). These findings hint at a mechanism whereby LEC afferents suppress recurrent activity through the activation of PV interneurons (Fig 6), suggesting a role that could switch responses to favour LOT over ASSN.

**Figure 6:**
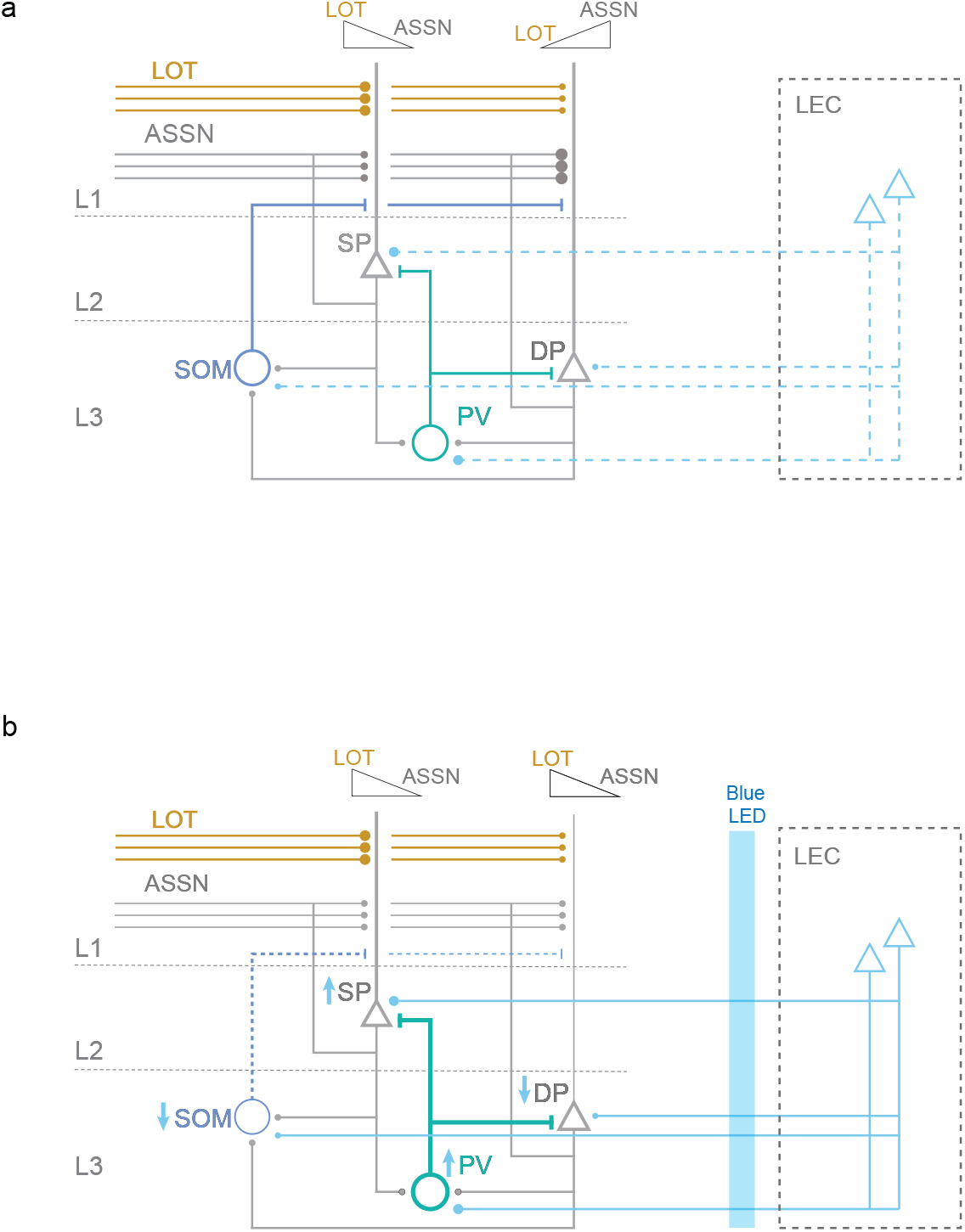
Reorganisation of PCx inputs induced by LEC activation. **A,** Sensory afferents arriving from the olfactory bulb through the LOT contact pyramidal neurons and also recruit ASSN circuit. SP neurons receive a larger fraction of LOT inputs than DP neurons. Conversely, DP neurons receive more inputs from ASSN compared with SP neurons. **B,** Optogenetic activation of excitatory LEC projections reconfigures PCx microcircuit. PV interneurons are are directly activated by LEC inputs, leading to a decrease in network recurrence. SOM interneurons are indirectly silenced by LEC activation disinhibiting apical dendrites of pyramidal neurons. This differential effect of LEC on PCx inhibitory circuits provokes an increase in pyramidal neurons’ response to LOT input while reducing their response to ASSN.

### LEC functional connectivity in the PCx microcircuit

Our recordings show that different classes of PCx neurons exhibited both EPSCs and IPSCs in response to LEC stimulation. However, neurons that were directly contacted (SP and PV neurons) displayed predominantly excitatory responses (a higher E/I balance) while neurons that responded primarily to the recurrent activity recruited by LEC, showed a predominant inhibitory component in their responses (DP and SOM, with a lower E/I balance). Heterogeneous distribution of afferent inputs has also been previously observed in the contacts that the Basolateral amygdala (BLA) establishes with the PCx (Luna & Morozov, 2012), where DP and PV neurons were preferentially contacted. The difference of both types of inputs arriving from higher processing areas suggest that each input differentially organises the activity in PCx, with BLA preferentially activating DP neurons and LEC preferentially activating SP neurons. However, both inputs produce strong activation of PV neurons, suggesting a common mechanism to inhibit recurrent activity.

### Inhibitory circuits recruited by LEC

Previous studies have demonstrated that lesions in the LEC of rats lead to an increased neuronal activity in the PCx at the population level (Chapuis et al., 2013). Based on this, it is argued that the LEC has a suppressive function on the PCx, consistent with an inhibition involving local inhibitory circuits. In this study, we were able to test this hypothesis and, furthermore, demonstrate that the inhibitory circuit involved is mediated by PV interneurons.

Although PV interneurons also receive inhibition, stimulation of LEC terminals was sufficient to trigger action potential firing from these neurons (Fig. S1). The activation of PV interneurons would have a negative effect on the recurrence of the circuit since these target principal neurons and control their spiking activity (Canto-Bustos et al., 2022; Large, Vogler, et al., 2016).

On the other hand, the silencing of SOM interneurons would promote a disinhibition at the level of dendrites in layer 1b. This would lead to enhanced integration of sensory afferents explaining the increase in the early spiking response of pyramidal neurons. It might also enable plasticity. It has been recently shown that SOM disinhibition facilitates long-term potentiation of the ASSN pathway after being associated with strong LOT stimulation (Canto-Bustos et al., 2022). The inputs that drive this disinhibition in physiological conditions are not known. We hypothesise that LEC afferents could be involved in such a mechanism, but further experiments should be carried out in order to prove this idea.

Overall, the differential impact of LEC on these two populations of inhibitory interneurons implies that its activation promotes a PCx microcircuit that is more sensitive to inputs from the LOT, while simultaneously reducing recurrence, thereby decreasing the response to this pathway.

### LEC effect in pyramidal neurons’ response to LOT

The temporal spiking pattern of neurons within PCx circuit plays a crucial role in neural dynamics. *In vivo*, global inhibition gives rise to narrow temporal windows of opportunity for generating an action potential (Poo & Isaacson, 2009). Neurons with slower activation kinetics are effectively silenced by inhibitory connections (Luna & Schoppa, 2008; Poo & Isaacson, 2009; Stokes & Isaacson, 2010; Suzuki & Bekkers, 2010). In our results of spiking latencies in response to stimulation of the LOT, both SP and DP neuronal populations displayed a dual-peaked distribution, with peaks around 10 ms and 20 ms, consistent with prior findings (Wiegand et al., 2011). We interpreted this bimodal distribution as arising from an early response to the afferent LOT pathway and a delayed response to recurrent activation of the ASSN pathway. This interpretation is supported by the observation that SP neurons, which exhibit strong responses to the afferent pathway, predominantly manifest an early response, while DP neurons, more sensitive to the ASSN pathway, display a pronounced delayed response.

Upon simultaneous stimulation of LOT and the LEC, we observed a temporal shift in the spiking distribution toward earlier responses. This shift was subtle for SP neurons, as they initially exhibited few late APs. However, for DP neurons, there was a notable decrease in late responses coupled with an increase in early spikes.

As previous studies have shown, single afferent stimulation is typically insufficient to elicit a spiking response in DP neurons (Large, Vogler, et al., 2016), indicating that input summation must be necessary to activate them. Changes in inhibition can have profound influence in determining the summation time window of neurons. In hippocampal pyramidal cells, weak inhibition at the dendritic level results in a broad integration window, while a strong perisomatic inhibition enforces precise coincidence detection in the soma (Pouille & Scanziani, 2001). Our results in DP neurons show that LEC stimulation narrows down the summation window. This suggests an increase in coincidence detection capabilities of this PCx neuronal population resulting in the locking of the response to the first afferent pulse, and the suppression of delayed APs.

Previous work has established that spatial information significantly influences PCx spiking activity (Federman et al., 2023; Mena et al., 2023; Poo et al., 2022). On the other hand, it is known that LEC plays a crucial role in the processing of the spatial location of objects (Beer et al., 2013; Chao et al., 2016; Deshmukh & Knierim, 2011; Hunsaker et al., 2013; Keene et al., 2016; Tsao et al., 2013; Van Cauter et al., 2013; D. I. G. Wilson, Langston, et al., 2013; D. I. G. Wilson, Watanabe, et al., 2013), the recognition of associative combinations of objects, places, and contexts with episodes (D. I. G. Wilson, Watanabe, et al., 2013), and in the association of attributes that make up an environment or local context (Kuruvilla & Ainge, 2017). This evidence, together with our results showing that LEC projections establish monosynaptic connections with PCx neurons and modulate the network activity, identify the LEC as a strong candidate to provide information about the spatial context to the representation of odours in the PCx.

The findings of this study demonstrate that LEC afferents exert distinct effects on the neuronal populations that compose the piriform cortex circuit. Activation of local inhibitory circuits mediated by PV interneurons leads to a dampening of recurrent activity. Conversely, inhibition of SOM interneurons results in dendritic disinhibition, facilitating enhanced integration of afferent signals. In essence, LEC activation induces a reorganisation of inputs within the circuit, favouring dominance of the LOT pathway over the ASSN pathway. These results shed light on the understanding of how contextual aspects of animal experience, in which LEC is involved, can influence odour processing potentially leading to a richer representation of odour objects.

## Acknowledgments

We thank members of the Marin-Burgin lab, the Refojo lab and Muraro lab for insightful discussions. We acknowledge International Development Research Centre IDRC108878 (AMB), Argentine Agency for the Promotion of Science and Technology, PICT2020-00360 (AMB), PICT 2020-1536 (NF), Swiss National Science Foundation (SNSF) SPIRIT 216044 (AMB) and FOCEM-Mercosur COF 03/11.

