## Supplementary figure for "Lateral entorhinal cortex afferents reconfigure the activity in piriform cortex circuits"

### Supplementary material

### Figure S1

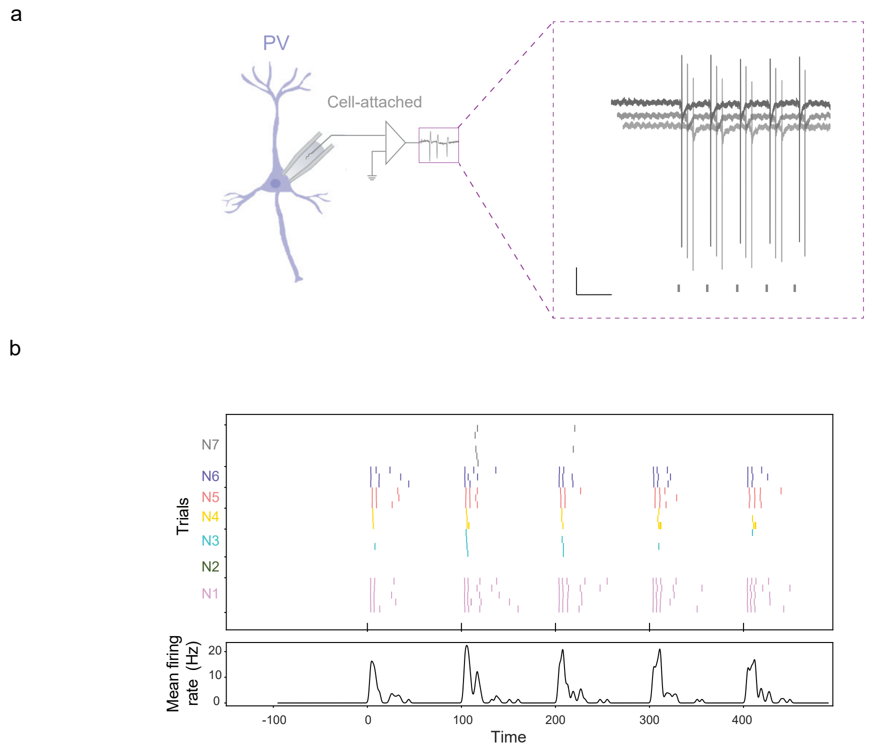


**Figure Supplementary 1: PV interneurons elicit a spiking response after LEC activation.**

**A,** Representative traces of a PV interneuron in *cell-attached* in response to a train of 5 pulses of blue light delivered at 10 Hz (each pulse is illustrated with a black line). Scale: x=125ms, y=50pA. **B,** TOP: Raster plot of PV interneurons spiking response. Each coloured line represents a single spike and each row a different sweep. Neurons are represented in different colours. Time zero corresponds to the first pulse of the train, and black lines on the x-axis indicate the time of each pulse. BOTTOM: Mean firing rate considering all neurons.
